# Regulatory non-coding small RNAs are diverse and abundant in an extremophilic microbial community

**DOI:** 10.1101/761684

**Authors:** Diego R. Gelsinger, Gherman Uritskiy, Rahul Reddy, Adam Munn, Katie Farney, Jocelyne DiRuggiero

## Abstract

Regulatory small RNAs (sRNAs) represent a major class of regulatory molecules that play large-scale and essential roles in many cellular processes across all domains of life. Microbial sRNAs have been primarily investigated in a few model organisms and little is known about the dynamics of sRNA synthesis in natural environments, and the roles of these short transcripts at the community level. Analyzing the metatranscriptome of a model extremophilic community inhabiting halite nodules (salt rocks) from the Atacama Desert with SnapT – a new sRNA annotation pipeline – we discovered hundreds of intergenic (itsRNAs) and antisense (asRNAs) sRNAs. The halite sRNAs were taxonomically diverse with the majority expressed by members of the *Halobacteria*. We found asRNAs with expression levels negatively correlated with that of their putative overlapping target, suggesting a potential gene regulatory mechanism. A number of itsRNAs were conserved and significantly differentially expressed (FDR<5%) between 2 sampling time points allowing for stable secondary structure modeling and target prediction. This work demonstrates that metatranscriptomic field experiments link environmental variation with changes in RNA pools and have the potential to provide new insights into environmental sensing and responses in natural microbial communities through non-coding RNA mediated gene regulation.

## INTRODUCTION

Non-coding RNAs (ncRNAs) are untranslated short transcripts that are found in the three domains of life where they play essential roles in many cellular processes (Gelsinger and DiRuggiero 2018b, Cech and Steitz 2014). In prokaryotes, a subset of these ncRNAs, thereby called small RNAs (sRNAs), are specifically involved in gene regulation through RNA-RNA mediated interactions, modulating core metabolic functions and stress related responses (Gottesman and Storz 2011). These sRNAs range from 50 to 500 nucleotides in size and can be of two types: *trans*-encoded sRNAs, also called intergenic sRNAs (itsRNAs), which bind their mRNA targets via imperfect base-pairing and can target multiple genes, including key transcription factors and regulators (Wagner and Romby 2015). itsRNAs can activate or inhibit translation initiation by interacting with the ribosome binding site (RBS) and/or modulating mRNA stability (Wagner and Romby 2015). In contrast, *cis*-encoded antisense RNAs (asRNAs) are transcribed on the DNA strand opposite their target gene and thus can act via extensive base pairing; they have been found to repress transposons and toxic protein synthesis (Wagner and Romby 2015).

The functional roles of microbial sRNAs have been extensively studied in a few model organisms and very little is known about the dynamics of sRNA synthesis in natural environments and the roles of these short transcripts at the community level (Carrier, Lalaouna, and Massé 2018, Gelsinger and DiRuggiero 2018b). To our knowledge, only two studies have reported the discovery of sRNAs in natural microbial communities (Shi, Tyson, and DeLong 2009, Bao et al. 2015). This paucity of knowledge suggests that an abundance of sRNAs remain to be discovered, in particular in extreme environments where they likely play essential roles in stress response (Clouet-d’Orval et al. 2018), inter-species communication, and/or cross-species RNA interference (Toyofuku, Nomura, and Eberl 2019, Cai et al. 2018, Tsatsaronis et al. 2018).

In hyper-arid deserts, microbial communities find refuge inside rocks as a survival strategy against the extreme conditions of their environment (Pointing and Belnap 2012). Such community inhabits halite (salt) nodules in Salars of the Atacama Desert, Chile, which is one of the oldest and driest deserts on Earth (Crits-Christoph et al. 2016, Finstad et al. 2017). The halite endolithic (within rock) community harbors mostly members of the Archaea (*Halobacteria*), unique *Cyanobacteria*, diverse heterotrophic bacteria, and a novel type of algae (Crits-Christoph et al. 2016, Finstad et al. 2017). The main source of liquid water for this community is from salt deliquescence (Davila et al. 2008) and it is sustained by CO_2_ fixed via photosynthesis (Crits-Christoph et al. 2016, Davila et al. 2015). While previous studies have demonstrated the role of sRNAs in the stress response of one of the members of this community, the halophilic archaeon *Haloferax volcanii* (Gelsinger and DiRuggiero 2018a, Kliemt, Jaschinski, and Soppa 2019), there is no information on any of the other members.

Here we used a combination of genome-resolved metagenomics and metatranscriptomics to investigate the role of sRNAs in the adaptive response of microorganisms inhabiting halite nodules. We developed an analytical pipeline, SnapT, built on our previous work on sRNAs with model organisms (Gelsinger and DiRuggiero 2018a), to enable the discovery of sRNAs at the community level. We found hundreds of sRNAs (both itsRNAs and asRNAs) in the halite community, including conserved sRNAs, validating our experimental approach. A number of itsRNAs were significantly differentially regulated between 2 sampling time points and, for a subset of these, we were able to perform structure and target prediction, deciphering their potential regulatory roles. Coupling metagenomics and metatranscriptomics with SnapT allows for the potential to uncover the complex regulatory networks that govern the state of a microbial community.

## MATERIAL AND METHODS

### Sample and weather data collection and nucleic acid extraction

Halite nodules were harvested in Salar Grande, an ancient evaporated lake in the Northern part of the Atacama Desert (Robinson et al. 2015) in February 2016 and 2017, 3 and 15 months after a major rain event (Uritskiy et al. 2019). All nodules were harvested within a 50m^2^ area as previously described (Robinson et al. 2015). The colonization zone of each nodule was grounded into a powder, pooling from 1-3 nodules until sufficient material was collected, and stored in the dark in dry conditions until DNA extraction in the lab. Samples used for RNA were stored in *RNAlater* at 4°C until RNA extraction in the lab. Genomic DNA was extracted as previously described (Robinson et al. 2015, Crits-Christoph et al. 2016) with the DNAeasy PowerSoil DNA extraction kit (QIAGEN). Total RNA was extracted from the fixed samples by first isolating the cells through gradual dissolving of the salt particles as previously described (Robinson et al. 2015, Crits-Christoph et al. 2016) and lysing them through mechanical bead beating with the RNAeasy PowerSoil RNA extraction kit (QIAGEN). Total RNA was then extracted from the lysate with a Quick-RNA miniprep kit (Zymo Research). RT-PCR was used to validate the absence of contaminating DNA in the total RNA used for RNA-seq libraries (Fig. S11).

### Library preparation

Whole genome sequencing libraries were prepared using the KAPA HyperPlus kit (Roche) as previously described (Uritskiy et al. 2019) and sequenced with paired 150bp reads on the HiSeq 2000 platform at the Johns Hopkins Genetic Resources Core Facility (GRCF). Total RNA-seq libraries were prepared with the SMARTer Stranded RNA-seq kit (Takara and Bell), using 25ng of RNA input and 12 cycles for library amplification. We sequenced 22 libraries from replicate samples from 2016 and 24 libraries from replicate samples from 2017.

### WMG sequence processing

The de-multiplexed WMG sequencing reads were processed with the complete metaWRAP v0.8.2 pipeline (Uritskiy, DiRuggiero, and Taylor 2018) with recommended databases on a UNIX cluster with 48 cores and 1024GB of RAM available. Detailed scripts for the entire analysis pipeline can be found at https://github.com/ursky/timeline_paper.

### SnapT for sRNA community identification

An analytic pipeline, SnapT for Small ncRNA Annotation Pipeline for (meta)Transcriptomic data, was adapted and developed from our previous work (Gelsinger and DiRuggiero 2018a) to find, annotate, and quantify intergenic and antisense sRNA transcripts from transcriptomic or metatranscriptomic data. Detailed scripts for the pipeline can be found at https://github.com/ursky/SnapT and search criteria were as follows: intergenic transcripts were at least 30 nt away from any gene or ORF on both strands; antisense transcripts were 30 nt away from any gene on their strand, but overlapped with a gene on the opposite strand by at least 10 nt; small peptides (<100 nt) were not counted as genes if they were encoded in a transcript that was more than 3 times their length; non-coding transcripts could not contain any reading frame greater than 1/3 of their lengths; predicted non-coding transcripts near contig edges were discarded and the minimum distance to the edge of a contig was dynamically computed such that the tips of contigs were not statistically enriched in annotated ncRNAs; small ncRNAs were between 50 nt and 500 nt in length; sRNA transcripts could not have significant homology with any protein in the NCBI_nr database (query cover>30%, Bitscore>50, evalue<0.0001, and identity>30%) and with any tRNA, RNase P, or signal recognition particle (SRP) model in the Rfam non-coding RNA database.

### Taxonomic assignment and distribution of sRNAs

The taxonomic origin of each annotated sRNA was taken to be as that of the contig on which it lay. The taxonomy of each contig was estimated by taking the weighted average of the taxonomic assignment of the genes encoded on it, as determined through the JGI IMG functional and taxonomic annotation service.

### Metatranscriptomic Correlation and Differential Expression Analysis

We used a read count-based differential expression analysis to identify differentially expressed sRNA and mRNA transcripts. The program featureCounts (Liao, Smyth, and Shi 2014) was used to rapidly count reads that map to the assembled RNA transcripts (described above) as previously described (Gelsinger and DiRuggiero 2018a). In order to account for organism abundance changes (as opposed to true transcript changes), we normalized the transcript read counts to the total read counts from the contig on which the transcript lies on. The read counts were then used in the R differential expression software package DESeq2 (Love, Huber, and Anders 2014) to calculate differential expression by determining the difference in read counts between 2016 normalized read counts from 2017 normalized read counts. The differentially expressed RNAs were filtered based on the statistical parameter of False Discovery Rate (FDR) and those that were equal to or under a FDR of 5% were classified as true differentially expressed transcripts. We carried out differential expression analysis using a pairwise Wald test to find any possible differences between years (Love, Huber, and Anders 2014). In parallel, normalized expression values were calculated using stringtie in transcripts per million (TPM). TPM of transcripts were normalized in the same way as read counts, except using contig TPM. TPM of transcripts was used for ranking of expression within samples as opposed to differential expression analysis.

### Regulatory element motif identification of sRNAs, structure and target prediction

50 nucleotides upstream from the sRNA transcript start coordinates were searched for transcription motifs (BRE and TATA-box for archaea and −35 and −10 consensus sequences for bacteria) using both multiple sequence alignments and visualization with WebLogo and motif searching with MEME (Gelsinger and DiRuggiero 2018a). Conserved sRNAs were identified using blastn against the NCBI nt database. Secondary structures of conserved sRNAs were predicted using sRNAs that had an e-value maximum of 1E-3, a sequence similarity of 70% or more, and 50% or more coverage with a NCBI nt database blastn hit; a minimum of 14 alignments were used in the program LocARNA using global alignment settings (Will et al. 2012). Lastly, putative targets were predicted for itsRNAs by searching for optimal sRNA-mRNA hybridization using the IntaRNA program with the no seed parameter (Mann, Wright, and Backofen 2017) and the reference genes for each respective MAG. Targets were ranked by lowest p-value. Expression levels for putative targets of antisense sRNAs were obtained from co-expression analysis of transcripts (Gelsinger and DiRuggiero 2018a). The sRNA and putative target mRNA TPM expression values were tracked across the replicates, and the Pearson correlation was computed.

### Enrichment cultures

Three types of culture medium were inoculated in triplicate with ~2 g of grounded halite colonization zones and incubated at 42°C with shaking at 220 rpm (Amerex Gyromax 737) for 1 to 2 weeks. Cells were harvested by centrifugation and nucleic acids extracted as described above. Media were: GN101 medium (Kish et al. 2009) containing 250 g of salt per L and 10 g of peptone as carbon source; Hv-YPC medium (Dyall-Smith 2009) containing 250 g of salt per L and 8.5 g of yeast extract, 1.7 g of peptone, and 1.7 of casamino acids as carbon sources; and IO containing 250 g of salt and the same carbon sources as the Hv-YPC medium. The taxonomic distribution of the cultures was obtained with 16S rRNA gene sequencing as previously described (Uritskiy et al. 2019).

#### sRNA validation

Total RNA extracted from environmental samples and enrichment cultures was converted into cDNA using the SuperScript III First-Strand Synthesis System (ThermoFisher). The cDNA was then amplified using primers designed for sRNAs identified in the halite community **(Table S1)**, as previously described (Meslier et al. 2018). Amplicons were sequenced using Sanger sequencing (GENEWIZ, South Plainfield, NJ).

#### Data availability

Raw sequencing data are available from the National Centre for Biotechnology Information under NCBI project ID PRJNA484015. The metagenome co-assembly and functional annotation are available from the JGI Genome Portal under IMG taxon OID 3300027982. Metatranscriptome data GEO # in process. Scripts for functional annotation, statistical analyses, differential expression, and figures are available at https://github.com/ursky/srna_metatranscriptome_paper.

## RESULTS

### Landscape of predicted sRNAs in the halite community and validation

We discovered hundreds of ncRNAs in an extremophilic community inhabiting halite nodules (salt rocks) in the Atacama Desert by using SnapT (https://github.com/ursky/SnapT), a pipeline adapted from our previous work on a model haloarchaeon present in the halite community (**Table 1; data S1**) (Gelsinger and DiRuggiero 2018a). We used metatranscriptomics data from multiple replicate samples collected in the field in 2016 and 2017 (21 and 24 replicates for 2016 and 2017, respectively; **Fig. S1**). Using SnapT, we aligned reads from stranded RNA-seq libraries to our reference co-assembled metagenome from a previous study (Uritskiy et al. 2019) and assembled the reads into transcripts (**Fig S2**). The transcripts were then intersected with the metagenome annotation as well as open reading frames to select for either novel transcripts on the opposite strand of coding transcripts (asRNAs) or for novel transcripts that fell into intergenic regions (itsRNAs). Putative ncRNA transcripts were then further enriched (**Fig. S2)** using a threshold at 5x and 10x assembly coverage in order to identify intergenic and antisense ncRNAs, respectively. (**Fig. S3; Table 1**). The size of these ncRNAs was then filtered from 50 to 500 nucleotides to produce a final set of non-coding sRNAs. The size distribution of these sRNAs was primarily between 50 and 200 nt for itsRNAs and above 200 nt for asRNAs. (**Fig. S4**).

**Table 1:** Summary of ncRNAs discovered in halite community

|  | **Number (%)<sup>\*</sup>** | **% in Archaea** | **% in Bacteria** |
| --- | --- | --- | --- |
| **Total ncRNAs** | 1538 (100) | 54 | 46 |
| **Rfam ncRNAs** | 79 (5) | 73 | 27 |
| **Conserved sRNAs<sup>\*\*</sup>** | 155 (10) | 60 | 40 |
| **Antisense sRNAs** | 925 (60) | 40 | 60 |
| **Intergenic sRNAs** | 613 (40) | 75 | 25 |

<sup>\*</sup>Percent from total ncRNAs
<sup>\*\*</sup> Conserved other than Rfam ncRNAs

The halite ncRNAs were taxonomically assigned to diverse members of the community; their distribution between Archaea (54%) and Bacteria (46%) (**Table 1**) was similar to that of the total metatranscriptomic reads for the community (**Fig. 1B** **and** **C**). In contrast, the taxonomic profile of the metagenome showed a larger contribution of bacterial reads and in particular of reads assigned to *Cyanobacteria* and *Bacteroidetes* (**Fig. 1A**). Because of the use of strand specific RNA-seq libraries, we could confidently identify both intergenic (it)sRNA, located between coding regions, and antisense (a)sRNA, overlapping with their putative target (**Table 1**). We found 3 times more itsRNAs in the Archaea than in the Bacteria, whereas asRNAs were more abundant in the Bacteria and more often associated with members of the *Cyanobacteria* (38%) and *Bacteriodetes* (15%) (**Table 1**; **Fig. 1D** **and** **E**). We also found 79 ncRNAs, that belong to 6 known families of RNAs present in the Rfam database (**Fig. S5**; **data S2**) (Kalvari et al. 2017), validating our experimental and computational approach. This database is a collection of RNA families, each represented by multiple sequence alignments, consensus secondary structures, and covariance models. Of the Rfam-conserved ncRNAs, 70% were assigned to archaea and included RNaseP RNAs, signal recognition particle RNAs (SRP RNAs), and tRNAs. Of the Rfam-conserved bacterial ncRNAs, most were from SRP RNAs and tRNA conserved families. In addition, a cobalamin riboswitch and the regulatory sRNA, CyVA-1, were detected in low abundance in the halite *Cyanobacteria*. We also found 3 ncRNAs (4%) from Eukarya, a tRNA, a U4 spliceosomal RNA, and a RNase for mitochondrial RNA processing (MRP). Using blastn analysis (max e-value of 1E-3, sequence similarity of 70% or more, coverage of 50% or more), we discovered another 155 ncRNAs that were conserved in the NCBI nt datasbase, with 60% from archaea and 40% from bacteria (**Table 1**). The majority were asRNAs (109), with only 44 itsRNAs. The conserved asRNAs most highly expressed (standardized tpm> 100) were all SPR RNAs in haloarchaea that were not found in the Rfam database. Of the conserved itsRNAs, we identified 3 tRNAs, 13 SRP RNAs, and 22 ncRNAs that were found in the genome of multiple species, all *Halobacteria*, but with no function assigned. The most highly expressed and conserved itsRNAs (standardized tpm> 100; 13 ncRNAs) were SRP RNAs not included in the Rfam database.

**Fig. 1.**
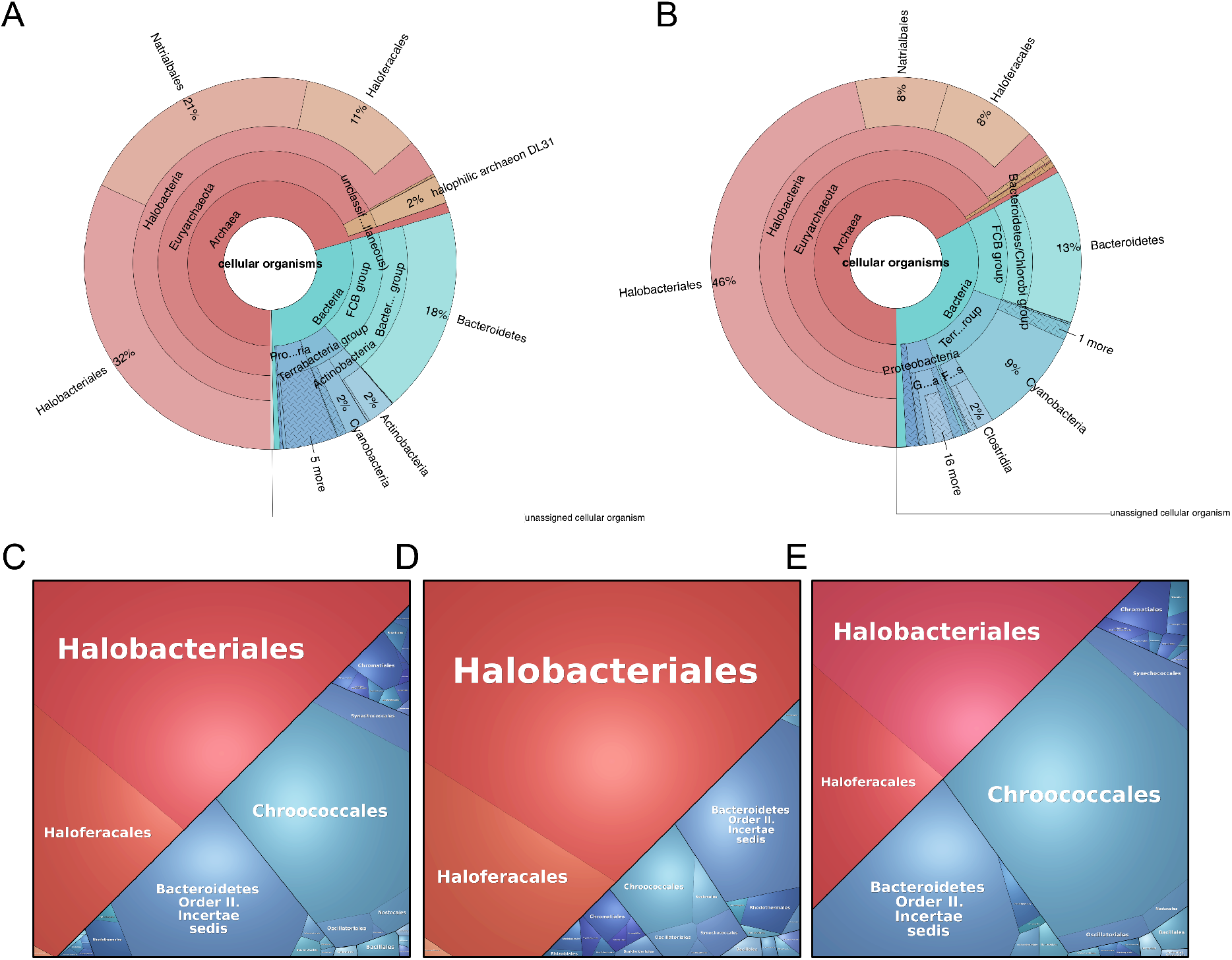
Taxonomic distribution. Krona graphs of (A) the halite metagenome based of DNA sequence reads and (B) the halite metatranscriptome based on RNA sequence reads; and Voronoi plots of (C) total sRNAs; (D) itsRNAs and (E) asRNAs discovered in the halite community.

Another validation of our findings was the presence of canonical promoter elements upstream of archaeal itsRNAs, suggesting that they were indeed *bona fide* transcripts that could recruit basal transcription factors (**Fig. S6**). We did not find significant promoter elements upstream of the bacterial itsRNAs, which might reflect the diversity of promoter elements across the various bacterial taxa we identified in the halite community. In contrast, no promoter elements were identified in the upstream regions of asRNA from both domains of life.

When looking at the expression levels of all itsRNAs normalized to contig abundances, we found that they were similar for both the 2016 and 2017 samples and slightly higher than that of the asRNAs, whereas the expression profile of the asRNAs was more variable across samples for both years (**Fig. S7**). Remarkably, the expression levels of itsRNAs and asRNAs for both years was 2-fold higher than that of protein encoding genes. Whereas there is an inherent bias in our approach to identify sRNAs at the community level (coverage threshold in SnapT) compared to protein encoding genes, this finding strongly indicates potential functional relevance for a number of these sRNAs.

We experimentally validated a number of sRNAs using RT-PCR with environmental and enrichment cultures (**Table S1**). Enrichments were performed with several media containing high (25%) and relatively low (18%) salt, and various combinations of carbon sources. Amplicon sequencing of the enrichments revealed that high salt and diverse carbon sources resulted in higher diversity of taxa, although haloarchaea dominated in all enrichments (**Fig. S8**). All validated sRNAs belong to haloarchaea with the exception of one from *Cyanobacteria*. Sequences of the PCR products confirmed that they were sRNAs and validated our computational approach.

### Relationship with target genes and putative function of community asRNAs

Using our strand-specific RNA-seq data, we were able to identify the overlap position of asRNAs to their antisense transcripts. We found that, in both Archaea and Bacteria, the majority of asRNAs start within the span of their cognate gene and end near the 5’ end of its mRNA. In both domains there is also an enrichment for asRNA-mRNA overlaps near the 5’ end of the mRNA. A similar trend has previously been reported in two species of archaea (Gelsinger and DiRuggiero 2018a, de Almeida et al. 2019).

We compared the expression level of asRNAs with that of their putative target genes and found that highly expressed asRNAs were associated with lowly express genes (**Fig. 2A**). Of gene pairs with asRNA expression >100 tpm and gene expression <0.1 tpm, most where from haloarchaea (77%), with 12% of *Cyanobacteria*, and 11% of other bacteria (*Bacteriodetes* and A*cinetobacter*) (**data S3**). Gene functions were enriched for transport (16%) and cell membrane/wall metabolism (5%), while most were hypothetical proteins (44%). Of the genes potentially negatively regulated by their cognate asRNAs, we found an archaeal regulator of the IclR family and potassium uptake protein TrkA. Only 2 asRNAs with high expression levels (>100 standardized tpm) were associated with genes with relatively high expression levels (>1 standardized tpm), while still being negatively correlated (Fig. 2A). The corresponding genes encoded for an iron complex outermembrane receptor protein from *Salinibacter* and a ABC-type sodium efflux pump permease subunit from a *Halobacteria*. When applying a stringent cut-off, we found 9 statistically significant and negatively correlated asRNA:gene pairs **(Fig. 2B)**. Four were from *Bacteroidetes*, 4 from *Halobacteria*, and 1 from an unidentified bacterium. At the functional level, transport systems, and in particular iron transport systems, were particularly enriched (**data S3**). In contrast, we did not find any significant positive regulation between asRNAs and their cognate genes. When adjusted for the carrying organism’s abundance, expressed as the average RNA read coverage of the contigs, we found that overall itsRNAs were more highly expressed than asRNAs (**Fig. 2C** **and** **2D**). Highly expressed sRNAs, for both types, were mostly carried by haloarchaea.

**Fig. 2.**
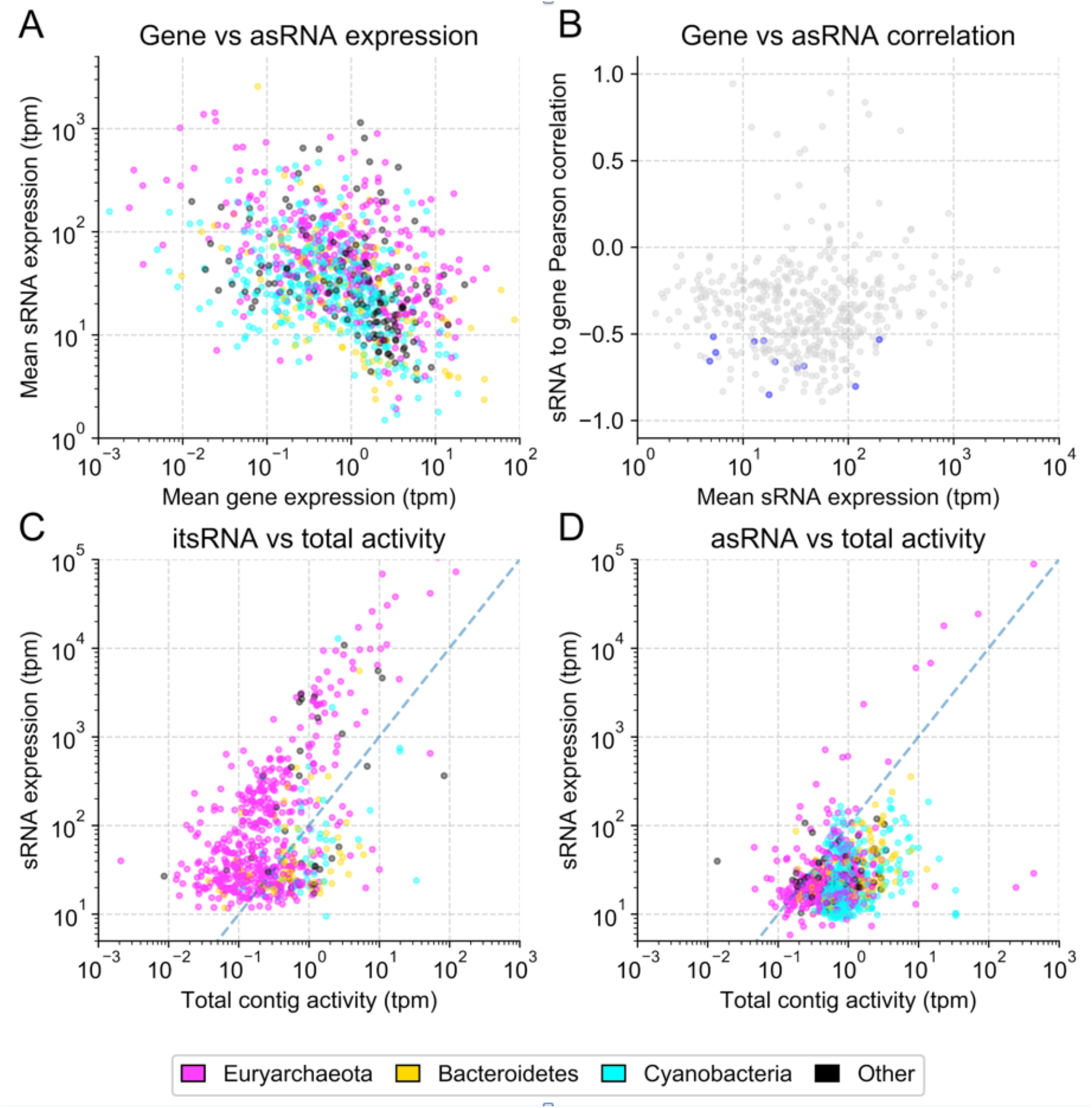
sRNA expression levels. (A) asRNAs and their putative targets mean expression levels (TPM); (B) Pearson correlations in expression level between asRNAs and their putative mRNA targets across the replicates, with significant correlations (pval<0.01) highlighted in blue; (C) average expression of itsRNA and average expression of (D) asRNAs over the average expression of the contigs on which they are found.

### Differential expression of itsRNAs at the community level and target prediction

Analysis of itsRNAs expression levels showed a clear separation between the 2016 and 2017 samples (**Fig. 3a**). We carried out a differential expression analysis and found that 109 (18%) of the regulatory itsRNAs were significantly differentially expressed (FDR<5%) between samples collected in 2016 and 2017 (**Fig. 3** **and data S4**), 3 and 15 months after a major rain event in the desert (Uritskiy et al. 2019). Of these, 72% were annotated as archaea and 28 % as bacteria and 16 were conserved in multiple genomes (14 from *Halobacteria* and 2 from *Cyanobacteria*). Conservation of differentially expressed itsRNAs allowed for structure modeling and, when high quality MAGs (>70% completion and <5% contamination) were available from the metagenome, target prediction (**Fig. 4** **and Fig. S10**). A number of non-differentially expressed itsRNAs were also conserved, providing additional opportunity for structure prediction; these included itsRNAs from *Halococcus* (STRG. 48671.1; 69 nt), *Halobellus limi* (STRG136887.1; 209 nt), and from a member of the *Nanohaloarchaea (*STRG.4577.1; 266 nt) (**Fig. S10A**).

**Fig. 3.**
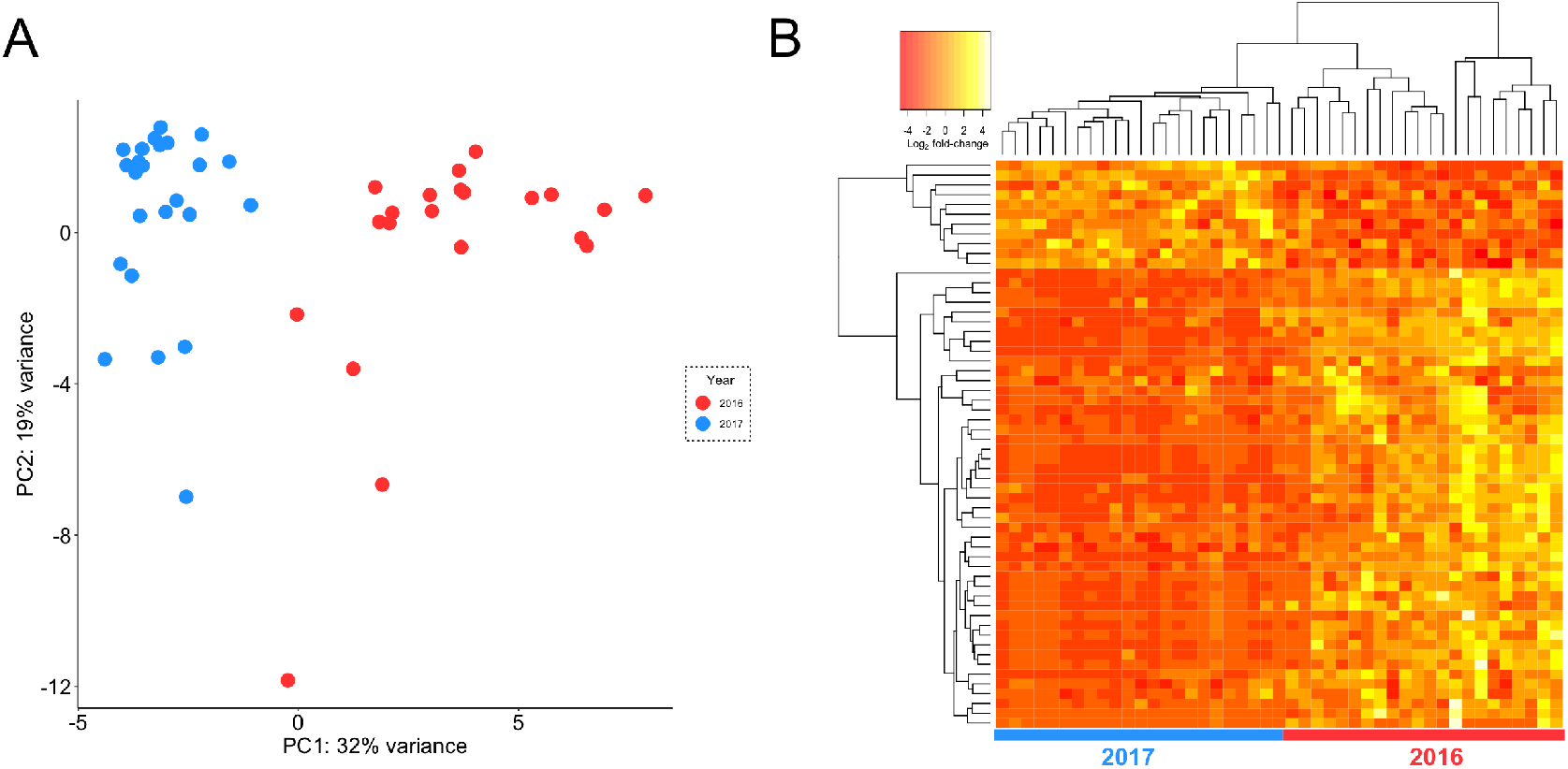
itsRNA differential expression. (A) PCA plot showing itsRNA expression levels clustered by year and (B) heat map of log_2_-transformed fold change for the top 50 significantly differentially expressed itsRNAs; each row is an itsRNA and each column a sample collected in 2016 or 2017.

**Fig. 4.**
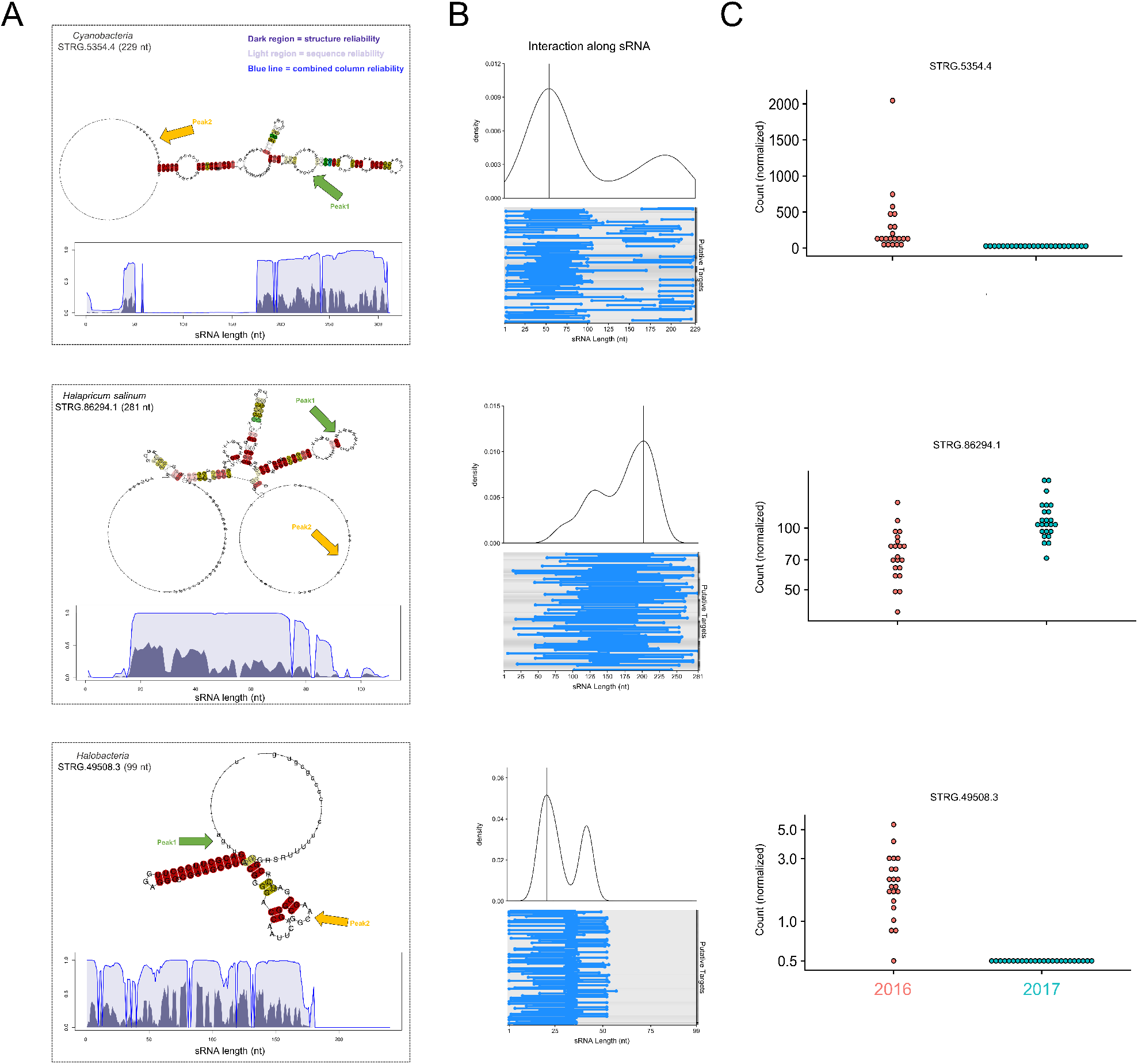
Predicted structure, target identification, and expression levels for selected differentially expressed itsRNAs. (A) 2D-layout of consensus structures with base pairs coloring showing sequence and structure conservation and interactions peaks (green and yellow arrows); STAR profile plots with dark regions indicating structure reliability, light regions representing sequence reliability, and thin lines showing the combined column-reliability as computed by LocARNA-P. (B) Interaction plots of itsRNAs and their predicted targets. The top graphs are density plots calculated from the top 100 putative targets, and on the bottom are dumbbell plots of interactions (blue dumbbells) along the length of the itsRNA for the top 100 predicted mRNA targets; interaction peaks are shown in green and yellow in the predicted structures; (C) Expression levels represented as normalized count for each itsRNA in 2016 and in 2017 across all samples.

All predicted structures displayed loop and stems regions that had high sequence conservation (light purple regions on sequence–structure-based alignment reliability [STAR] profile plots) and high structure conservation (dark purple), and line plots representing the reliability of the predictions as calculated by LocaRNA (**Fig. 4** **and Fig. S10B**). Density plots combined with dumbbell plots were used for visualizing predicted interactions between itsRNAs and their putative targets, using IntaRNA data from the top 100 most reliable interaction predictions with the lowest free energy of hybridization (Mann, Wright, and Backofen 2017) **(Fig. 4)**. High confidence assignments were obtained for 4 differentially expressed itsRNAs from *Cyanobacteria*, *Halapricum salinun*, and a member of the *Halobacteria* (**data S5**) More than one interaction peak were derived from density plots; peak 1 (green) corresponded to the highest interaction density, which mapped to loop regions in the itsRNA secondary structure with high sequence and structure conservation, respectively, and was thus a confident assignment as an interaction region, whereas Peak 2 (yellow) was a less confident assignment structurally despite high interaction density (**Fig. 4** **and Fig. S10B**).

Using this information, we identified the most probable targets for *Cyanobacteria* STRG.5354.4 candidate itsRNA (229 nt). This itsRNA was conserved as a 6S regulatory RNA in the rfam database, which in bacteria is found to inhibit transcription by binding directly to the housekeeping holoenzyme form of RNA polymerase (Wassarman 2018). Of the top 50 most probable targets for STRG.5354.4, which were those with the lowest free energy of hybridization between itsRNA and targets, were cation:H+ antiporters [shown to be involved in osmoregulation (Krulwich, Hicks, and Ito 2009)], a PleD family two-component response regulator, the photosystem I PsaB protein, chemotaxis transducers, and proteins involved in energy metabolism. Most probable targets for differentially expressed itsRNA, STRG. 86294.1 (281 nt) from *Halapricum salinum* included various transporters and putative membrane and cell wall associated proteins; notably an ammonium transporter (Amt family), an alkanesulfonate monooxygenase SsuD from a gene cluster expressed under sulfate or cysteine starvation (Eichhorn, van der Ploeg, and Leisinger 1999), and several proteins involved in cofactors and vitamin metabolism. Predicted targets with the lowest free energy of hybridization for STRG.49508.3 candidate itsRNA (99 nt) from *Halobacteria* were elongation factor 1-alpha, which promotes the GTP-dependent binding of aminoacyl-tRNA to the A-site of ribosomes during protein biosynthesis, several ribosomal proteins, and a number of hypothetical proteins. Target prediction for *Cyanobacteria* STRG.5356.1 candidate itsRNA (242 nt) included molecular chaperones (DnaK and DnaJ classes), a cell division protease FtsH, and a number of uncharacterized proteins.

## DISCUSSION

The roles of regulatory sRNAs have been extensively studie<u>d</u> in bacterial, and to a lesser extent, in archaeal model systems (Carrier, Lalaouna, and Massé 2018, Gelsinger and DiRuggiero 2018b) but, to date, only two studies have reported the discovery of sRNAs in microbial communities. <u>I</u>n one study, Shi *et al*. (Shi, Tyson, and DeLong 2009) used metatranscriptomic data to identify unique microbial sRNAs in the ocean’s water column while the study by Bao et al. (Bao et al. 2015) revealed extensive antisense transcription in the human gut microbiota, also using metatranscriptomic datasets. Efforts have also been made to mine publically available databases for sRNA discovery (Weinberg et al. 2017) but this was still addressing the role of sRNAs in single microorganisms.

One major difficulty in obtaining metatranscriptomic data from natural microbial communities, in particular from extreme environments, is the low amount of biomass that can be collected, resulting in low RNA yields (Uritskiy and DiRuggiero 2019). This, in turn, prevents attempts at ribo-depletion, resulting in a decreased number of non-ribosomal RNA reads available for analysis. Nevertheless, using SnapT, a flexible pipeline to process metagenomics and metatranscriptomic data, we report the discovery of hundreds of diverse sRNAs from an extremophilic community inhabiting halite nodules in the Atacama Desert. In the process, we applied extensive quality control with coverage thresholding, correction for contig edge mis-annotation, and the removal of potential non-ncRNAs through sequence and homology searches. While this approach might potentially result in false negatives, and may bias our findings toward the most highly expressed sRNAs in the community, it also insured the robustness of our sRNA predictions by minimizing the number false positives. The identification of ncRNAs in the halite community that belong to the Rfam database **(Kalvari et al. 2017)**, together with experimental validation of a number of sRNAs with environmental and enrichment cultures, substantiated our analytical approach. Additionally, expression levels of sRNAs 2-fold higher than that of protein encoding genes, strongly indicates potential functional relevance for a number of these sRNAs.

The taxonomic composition of the halite sRNAs matched that of the community’s metatranscriptomic profile, reflecting the contribution of the most active members, including *Cyanobacteria*, *Bacteriodetes*, and a number of *Halobacteria*. We found significantly more itsRNAs in the archaea than in the bacteria and the trend was reverse for the asRNAs. This novel finding is representative of published work in model organisms where a wide range of sRNAs has been found so far in prokaryotes, from less than a dozen to more than a thousand per genome (Carrier, Lalaouna, and Massé 2018, Gelsinger and DiRuggiero 2018b).

Antisense sRNAs overlap their putative targets providing insights into their functional role (Wagner and Romby 2015). In the halite community, we found that asRNAs expression levels were negatively correlated with that of their putative targets, with highly expressed asRNAs overlapping lowly expressed protein encoding genes. A similar trends was reported in the haloarchaeon *Haloferax volcanii*, when investigating oxidative stress responsive sRNAs, and most of the putative targets were transposase genes (Gelsinger and DiRuggiero 2018a). Putative target gene functions in our study were mostly from haloarchaea and enriched for transport systems, cell membrane and cell wall metabolism, with a large number of hypotheticals. Of particular interest, was an archaeal IcIR transcription regulator; these regulators are known to be involved in diverse physiological functions, including multidrug resistance, degradation of aromatics, and secondary metabolites production (Molina-Henares et al. 2006) and are distributed in a wide range of prokaryotes, including Archaea (Perez-Rueda et al. 2018). Also of interest, was a Trk potassium uptake system, also found in both bacteria and archaea and essential for the maintenance of high intracellular potassium in salt-in strategists (Oren 2013). In contrast, we did not find any significant positive regulation between asRNAs and their cognate genes, which might be due to the inherent quality of our data set, i.e. no ribo-depletion and heterogeneity across replicates (Uritskiy and DiRuggiero 2019). Alternatively, it might also reflect promiscuous transcription processes as argued when considering the functionality of asRNAs (Lloréns-Rico et al. 2016). Other arguments in favor of spurious transcription was the size distribution for asRNAs found in the halite community, which was significantly larger than that of itsRNAs, low expression level when adjusted for organism abundance when compared to itsRNAs, and the absence of canonical regulatory elements in the upstream regions of asRNAs. However, we found also putative target functions that reflected the environmental challenges faced by members of this extremophile community, such as osmoregulation and nutrient uptake, indicating that these asRNAs might indeed regulate fundamental biological functions at the community level.

We previously showed that the halite community dramatically shifted its taxonomic and functional composition after a major rain event in 2015, and while it recovered at the functional level in 2017, 15 months after the rain, members of the communities were permanently replaced (Uritskiy et al. 2019). Here we found that 18% of the halite community itsRNAs were significantly differentially expressed (FDR <5%) between samples collected in 2016 and 2017 (3 and 15 months after the rain, respectively), potentially indicating a transcriptional response to changes in environmental conditions. Intergenic sRNAs are of particular interest because they can target multiple genes, including key transcription factors and regulators (Gottesman and Storz 2011). As a consequence, a single sRNA can modulate the expression of large regulons and thus have a significant effect on metabolic processes (Carrier, Lalaouna, and Massé 2018). However, they do not overlap their target genes or bind their targets mRNAs with perfect complementary, which make finding targets for these sRNAs very challenging without genetic tools (Gelsinger and DiRuggiero 2018b).

To solve this problem at the community level, we focused on itsRNAs that were conserved and for which we could perform structural prediction. The intersection of this small subset of sRNAs with high quality MAGs that could be used as reference genomes, yielded confident target predictions for 4 differentially expressed itsRNAs, giving insights into metabolic functions potentially regulated by sRNAs at the community level. These included transporters, particularly related to osmotic stress, nutrient uptake and starvation, and pathways for chemotaxis and energy production and conversion. These pathways reflect the environmental challenges members of the halite communities are subjected to, including osmotic adjustments to climate perturbation (Uritskiy et al. 2019) and competition for nutrients in a near-close system with primary production as the major source of organic carbon (Crits-Christoph et al. 2016). Using the genomic context of sRNAs from the ocean’s water column microbial communities, Shi *et al*. (Shi, Tyson, and DeLong 2009) reported similar metabolic functions, underlying specific regulatory needs for natural communities. In contrast, genes with antisense transcription to asRNAs identified in the human gut microbiome were mostly transposase genes with a small component of bacterial house-keeping genes (Bao et al. 2015). It important to note that no computational target prediction, using sRNA conserved predicted structure, was reported in either study.

Regulation of transcription by 6S sRNA has been shown to increase competitiveness and long-term survival in bacteria (Wassarman 2018), suggesting an important role for *Cyanobacteria* candidate sRNA STRG.5354.4, identified as a 6S sRNA. Because of high RNA-seq coverage of the *Cyanobacteria* MAGs, we could show that 40% of the top 50 targets for sRNA STRG.5354.4 were differentially regulated and more highly expressed in 2016, suggesting positive regulation by this sRNAs onto its putative targets. Transcriptional factors and regulators were also found as putative targets of differentially regulated itsRNAs in the halite community, underlying the capacity of microbial sRNAs to modulate the expression of large regulons (Gottesman and Storz 2011, Gelsinger and DiRuggiero 2018b, Nitzan, Rehani, and Margalit 2017). Finally, a candidate itsRNAs from the *Halobacteria* had a number of predicted targets associated with ribosomal proteins and proteins involved in translation processes. This finding, together with a recent study in *H. volcanii* **(Wyss et al. 2018)**, support the idea of sRNA modulation of protein biosynthesis in the Archaea. A potential framework for mechanisms for sRNA regulation of translation might be provided by a report, in the haloarchaeon *Halobacterium salinarum*, of modular translation subsystems that might selectively translate a subset of the transcriptome under specific growth conditions **(Raman et al. 2018)**.

### Conclusion

In this study, we characterized the taxonomic and functional landscape of sRNAs across two domains of life in an extremophilic microbial community, demonstrating that asRNAs and itsRNAs can be reliably identified from natural environmental communities. To facilitate this work, we built a flexible pipeline, SnapT (https://github.com/ursky/SnapT), leveraged by our expertise of sRNA biology in a model halophilic archaeon, and which is available to use with metatranscriptomic data from any community. We demonstrated that we could perform target prediction and correlate expression levels between itsRNAs and predicted target mRNAs, paving the way for novel discoveries that have never been done at the community level. While additional work with enrichment cultures remain to be done to fully characterize the functional roles of sRNAs from the halite community, and their mechanism of action, these differentially expressed sRNAs for which we found putative targets show the power of community-level, culture-independent approach analysis for gene regulation processes.

## Author Contributions

JDR design the study, collected field samples, and wrote the manuscript. DRG and GU collected field samples, conducted experiments, analyzed the data, and edited the manuscript. RR and AM performed experimental target validation. KF grew and characterized enrichment cultures. All authors approved the final version for submission.

## Acknowledgments

This work was supported by NASA grant 18-EXO18-0091.

## Conflict of Interest

The authors declare that they have no conflict of interest.

## References

Bao, G. H., M. J. Wang, T. G. Doak, and Y. Z. Ye. 2015. “Strand-specific community RNA-seq reveals prevalent and dynamic antisense transcription in human gut microbiota.” Front Microbiol 6. doi: ARTN 89610.3389/fmicb.2015.00896.

Cai, Qiang, Lulu Qiao, Ming Wang, Baoye He, Feng-Mao Lin, Jared Palmquist, Sienna-Da Huang, and Hailing Jin. 2018. “Plants send small RNAs in extracellular vesicles to fungal pathogen to silence virulence genes.” Science 360 (6393):1126. doi: 10.1126/science.aar4142.

Carrier, Marie-Claude, David Lalaouna, and Eric Massé. 2018. “Broadening the Definition of Bacterial Small RNAs: Characteristics and Mechanisms of Action.” Ann Rev Microbiol 72 (1):141–161. doi: 10.1146/annurev-micro-090817-062607.

Cech, Thomas R, and Joan A Steitz. 2014. “The Noncoding RNA Revolution-Trashing Old Rules to Forge New Ones.” Cell 157 (1):77–94. doi: 10.1016/j.cell.2014.03.008.

Clouet-d’Orval, Béatrice, Manon Batista, Marie Bouvier, Yves Quentin, Gwennaele Fichant, Anita Marchfelder, and Lisa-Katharina Maier. 2018. “Insights into RNA-processing pathways and associated RNA-degrading enzymes in Archaea.” FEMS Microbiol Rev 42 (5):579–613. doi: 10.1093/femsre/fuy016.

Crits-Christoph, A., D.R. Gelsinger, B. Ma, J. Wierzchos, J. Ravel, C. Ascaso, O. Artieda, A. Davila, and J. DiRuggiero. 2016. “Functional analysis of the archaea, bacteria, and viruses from a halite endolithic microbial community.” Env. Microbiol. 18:2064–2077. doi: 10.1111/1462-2920.13259.

Davila, A.F., B. Gomez-Silva, A. de los Rios, C. Ascaso, H. Olivares, C.P. McKay, and J. Wierzchos. 2008. “Facilitation of endolithic microbial survival in the hyperarid core of the Atacama Desert by mineral deliquescence.” J. Geophys. Res. 113 (G01028):G01028, doi:10.1029/2007JG000561.

Davila, A.F., I. Hawes, J. Garcia, D.R. Gelsinger, J. DiRuggiero, C. Ascaso, A. Osano, and J. Wierzchos. 2015. “In situ metabolism in halite endolithic microbial communities of the hyperarid Atacama Desert.” Front Microbiol http://dx.doi.org/10.3389/fmicb.2015.01035.

de Almeida, João Paulo Pereira, Ricardo Z. N. Vêncio, Alan P. R. Lorenzetti, Felipe Ten Caten, José Vicente Gomes-Filho, and Tie Koide. 2019. “The Primary Antisense Transcriptome of *Halobacterium salinarum* NRC-1.” Genes 10 (4):280. doi: 10.3390/genes10040280.

Dyall-Smith, M. 2009. “The Halohandbook – Protocols for haloarchaeal genetics.” *Available at* https://www.researchgate.net/publication/278741334_The_Halohandbook_v73

Eichhorn, Eric, Jan R. van der Ploeg, and Thomas Leisinger. 1999. “Characterization of a Two-component Alkanesulfonate Monooxygenase from *Escherichia coli.*” J Biol l Chem 274 (38):26639–26646. DOI:10.1074/jbc.274.38.26639

Finstad, K. M., A. J. Probst, B. C. Thomas, G. L. Andersen, C. Demergasso, A. Echeverria, R. G. Amundson, and J. F. Banfield. 2017. “Microbial Community Structure and the Persistence of Cyanobacterial Populations in Salt Crusts of the Hyperarid Atacama Desert from Genome-Resolved Metagenomics.” Front Microbiol 8. doi: ARTN 143510.3389/fmicb.2017.01435.

Gelsinger, D. R., and J. DiRuggiero. 2018a. “Transcriptional Landscape and Regulatory Roles of Small Noncoding RNAs in the Oxidative Stress Response of the Haloarchaeon Haloferax volcanii.” J Bacteriol 200 (9). doi: ARTN e00779-1710.1128/JB.00779-17.

Gelsinger, Diego, and Jocelyne DiRuggiero. 2018b. “The Non-Coding Regulatory RNA Revolution in Archaea.” Genes 9 (3):141. doi: 10.3390/genes9030141.

Gottesman, S., and G. Storz. 2011. “Bacterial small RNA regulators: versatile roles and rapidly evolving variations.” Cold Spring Harb Perspect Biol 3 (12). doi: 10.1101/cshperspect.a003798.

Kalvari, Ioanna, Joanna Argasinska, Natalia Quinones-Olvera, Eric P. Nawrocki, Elena Rivas, Sean R. Eddy, Alex Bateman, Robert D. Finn, and Anton I. Petrov. 2017. “Rfam 13.0: shifting to a genome-centric resource for non-coding RNA families.” NAR 46 (D1):D335–D342. doi: 10.1093/nar/gkx1038.

Kish, A., G. Kirkali, C. Robinson, R. Rosenblatt, P. Jaruga, M. Dizdaroglu, and J. DiRuggiero. 2009. “Salt shield: intracellular salts provide cellular protection against ionizing radiation in the halophilic archaeon, Halobacterium salinarum NRC-1.” Environ Microbiol 11 (5):1066. DOI: 10.1111/j.1462-2920.2008.01828.x

Kliemt, Jana, Katharina Jaschinski, and Jörg Soppa. 2019. “A Haloarchaeal Small Regulatory RNA (sRNA) Is Essential for Rapid Adaptation to Phosphate Starvation Conditions.” Front Microbiol 10:1219–1219. doi: 10.3389/fmicb.2019.01219.

Krulwich, Terry A., David B. Hicks, and Masahiro Ito. 2009. “Cation/proton antiporter complements of bacteria: why so large and diverse?” Mol Microbiol 74 (2):257–260. doi: 10.1111/j.1365-2958.2009.06842.x.

Liao, Yang, Gordon K. Smyth, and Wei Shi. 2014. “featureCounts: an efficient general purpose program for assigning sequence reads to genomic features.” Bioinformatics 30 (7):923–930. doi: 10.1093/bioinformatics/btt656.

Lloréns-Rico, Verónica, Jaime Cano, Tjerko Kamminga, Rosario Gil, Amparo Latorre, Wei-Hua Chen, Peer Bork, John I. Glass, Luis Serrano, and Maria Lluch-Senar. 2016. “Bacterial antisense RNAs are mainly the product of transcriptional noise.” Science adv 2 (3):e1501363–e1501363. doi: 10.1126/sciadv.1501363.

Love, M. I., W. Huber, and S. Anders. 2014. “Moderated estimation of fold change and dispersion for RNA-seq data with DESeq2.” Genome Biol 15 (12):550. doi: 10.1186/s13059-014-0550-8.

Mann, Martin, Patrick R. Wright, and Rolf Backofen. 2017. “IntaRNA 2.0: enhanced and customizable prediction of RNA–RNA interactions.” NAR 45 (W1):W435–W439. doi: 10.1093/nar/gkx279.

Meslier, V., M.C. Casero, M. Daily, J. Wierchos, C. Ascaso, O. Artieda, P.R. McCullough, and J. DiRuggiero. 2018. “Fundamental drivers for endolithic microbial community assemblies in the hyperarid Atacama Desert.” Env. Microbiol. 20:1765–1781. https://doi.org/10.1111/1462-2920.14106.

Molina-Henares, Antonio J., Tino Krell, Maria Eugenia Guazzaroni, Ana Segura, and Juan L. Ramos. 2006. “Members of the IclR family of bacterial transcriptional regulators function as activators and/or repressors.” FEMS Microbiol Rev 30 (2):157–186. doi: 10.1111/j.1574-6976.2005.00008.x.

Nitzan, Mor, Rotem Rehani, and Hanah Margalit. 2017. “Integration of Bacterial Small RNAs in Regulatory Networks.” Ann Rev Biophys 46 (1):131–148. doi: 10.1146/annurev-biophys-070816-034058.

Oren, Aharon. 2013. “Life at high salt concentrations, intracellular KCl concentrations, and acidic proteomes.” Front Microbiol 4:315–315. doi: 10.3389/fmicb.2013.00315.

Perez-Rueda, Ernesto, Rafael Hernandez-Guerrero, Mario Alberto Martinez-Nuñez, Dagoberto Armenta-Medina, Israel Sanchez, and J. Antonio Ibarra. 2018. “Abundance, diversity and domain architecture variability in prokaryotic DNA-binding transcription factors.” PloS one 13 (4):e0195332–e0195332. doi: 10.1371/journal.pone.0195332.

Pointing, Stephen B., and Jayne Belnap. 2012. “Microbial colonization and controls in dryland systems.” Nature Rev Microbiol 10:551. doi: 10.1038/nrmicro2831.

Raman, A.V., A.n López García de Lomana, U. Kusebauch, M. Pan, S. Turkarslan, R. L. Moritz, and N. S. Baliga. 2018. “Context-Specific Regulation of Coupled Transcription-Translation Modules Predicts Pervasive Ribosome Specialization.” *Available at SSRN:* http://dx.doi.org/10.2139/ssrn.3155765

Robinson, C. K., J. Wierzchos, C. Black, A. Crits-Christoph, B. Ma, J. Ravel, C. Ascaso, O. Artieda, S. Valea, M. Roldan, B. Gomez-Silva, and J. DiRuggiero. 2015. “Microbial diversity and the presence of algae in halite endolithic communities are correlated to atmospheric moisture in the hyper-arid zone of the Atacama Desert.” Environ Microbiol 17:299–315. doi: 10.1111/1462-2920.12364.

Shi, Y. M., G. W. Tyson, and E. F. DeLong. 2009. “Metatranscriptomics reveals unique microbial small RNAs in the ocean’s water column.” Nature 459 (7244):266–U154. doi: 10.1038/nature08055.

Takara, T. J., and S. P. Bell. 2009. “Putting two heads together to unwind DNA.” Cell 139 (4):652–654. doi: 10.1016/j.cell.2009.10.037.

Toyofuku, Masanori, Nobuhiko Nomura, and Leo Eberl. 2019. “Types and origins of bacterial membrane vesicles.” Nature Rev Microbiol 17 (1):13–24. doi: 10.1038/s41579-018-0112-2.

Tsatsaronis, James A., Sandra Franch-Arroyo, Ulrike Resch, and Emmanuelle Charpentier. 2018. “Extracellular Vesicle RNA: A Universal Mediator of Microbial Communication?” Trends Microbiol 26 (5):401–410. doi: 10.1016/j.tim.2018.02.009.

Uritskiy, G.V., and J. DiRuggiero. 2019. “Applying Genome-Resolved Metagenomics to Deconvolute the Halophilic Microbiome.” Gene 10:220. doi: 10.3390/genes10030220.

Uritskiy, Gherman, Samantha Getsin, Adam Munn, Benito Gomez-Silva, Alfonso Davila, Brian Glass, James Taylor, and Jocelyne DiRuggiero. 2019. “Halophilic microbial community compositional shift after a rare rainfall in the Atacama Desert.” ISMEJ doi: 10.1038/s41396-019-0468-y.

Uritskiy, Gherman V., Jocelyne DiRuggiero, and James Taylor. 2018. “MetaWRAP—a flexible pipeline for genome-resolved metagenomic data analysis.” Microbiome 6 (1):158. doi: 10.1186/s40168-018-0541-1.

Wagner, E. Gerhart H., and Pascale Romby. 2015. “Chapter Three - Small RNAs in Bacteria and Archaea: Who They Are, What They Do, and How They Do It.” In Advances in Genetics, edited by Theodore Friedmann, Jay C. Dunlap and Stephen F. Goodwin, 133–208. Academic Press.

Wassarman, Karen M. 2018. “6S RNA, a Global Regulator of Transcription.” Microbiol Spectr 6 (3). doi: 10.1128/microbiolspec.RWR-0019-2018.

Weinberg, Zasha, Christina E. Lünse, Keith A. Corbino, Tyler D. Ames, James W. Nelson, Adam Roth, Kevin R. Perkins, Madeline E. Sherlock, and Ronald R. Breaker. 2017. “Detection of 224 candidate structured RNAs by comparative analysis of specific subsets of intergenic regions.” NAR 45 (18):10811–10823. doi: 10.1093/nar/gkx699.

Will, Sebastian, Tejal Joshi, Ivo L. Hofacker, Peter F. Stadler, and Rolf Backofen. 2012. “LocARNA-P: Accurate boundary prediction and improved detection of structural RNAs.” RNA 18 (5):900–914. doi: 10.1261/rna.029041.111.

Wyss, Leander, Melanie Waser, Jennifer Gebetsberger, Marek Zywicki, and Norbert Polacek. 2018. “mRNA-specific translation regulation by a ribosome-associated ncRNA in Haloferax volcanii.” Scientific Rep 8 (1):12502. doi: 10.1038/s41598-018-30332-w.

