## supplementary material for "Regulatory non-coding small RNAs are diverse and abundant in an extremophilic microbial community"

### Supplementary Materials

**Table of content:**

Table S1 Experimentally validated itsRNAs

Fig. S1 Experimental design
Fig. S2 Flow chart for SnapT methodology
Fig. S3 Thresholding for its and asRNAs
Fig. S4 Length distribution of itsRNAs and asRNAs
Fig. S5 Rfam-conserved sRNAs indentified the halite community
Fig. S6 Regulatory regions for asRNAs and itsRNAs and for archaea and bacteria
Fig. S7 Expression level of itsRNAs, asRNAs, and protein encoding genes
Fig. S8 Taxonomy distribution of halite enrichment cultures
Fig. S9 asRNA overlap distribution with their putative mRNAs
Fig. S10 Additional structure and target predictions
Fig. S11 RT-PCR

**Additional data files:**

data S1. sRNAs features, expression levels, and conserved
data S2. Rfam ncRNAs
data S3: asRNA; gene pairs with TPM
data S4. Significant differential expressed sRNAs
data S5: IntaRNA interactions for itsRNAs

**Table S1:** Experimentally validated isRNAs

| | | | RT-PCR amplicon with | | |
|---|---|---|---|---|---|
| sRNA_ID | Taxonomy | Primer set | Environ. sample | Enrichment cultures | Sequence |
| STRG.134829.3 | *Halothece* | | 17A2, 9B2 | IO-YPC; GN101; Hv-YPC | > STRG.134829.3<br>AACCTTACGGTTTGTCCAAAT<br>TAACCTCTGTACCGGTAGTCC<br>ATCGGAAATTGACGAGCAAC<br>CCTGAATTGACAGGGTGTCC<br>AATTAAACTGGATGGAGATA<br>CGATGACTAGCCGTTAGGTG<br>GTCAGCCTGCTAACCCTCGAT<br>TTGAGTGGCGGAGAACCTGA<br>GTGATTAGGTTCTAGAATCTC<br>CCACCATAATCTTTGATTTGG<br>TGGTGAGAGT |
| STRG.66426.1 | *Salinarchaeum* | | 17B2 | IO-YPC; GN101; Hv-YPC | > STRG.66426.1<br>GGTCGGACTAGGCTGGGCGG<br>TTAGGCCCCGCTCCGACGCC<br>CGCAGTACGGTCTTCAGCGG<br>GGGCCGAACCCGGGGACGTC<br>CGGTCAGACCGGGACGGGCC<br>TCGGAAGCCAACGTCGAAGC<br>CTCGTCCCTCGGGGACGACG<br>GTCCACGGCGGTCGCCTGCA<br>GGGGCGCGTTGTCGTGGTTC<br>GTCGGCGGCACCGGGTCAGG<br>CGCGGAAGCGAGCAGCCCAC<br>CGTCG |
| STRG.104813.1 | *Halomicrobium* | | none | Hv-YPC; GN101 | > STRG.104813.1<br>GGACTCCAGTTTCAGGCCGT<br>GAAACCGCCGTTAGTGCGAT<br>GTAGCGCCGAAAACAATCAC<br>AACAATCACTAT |

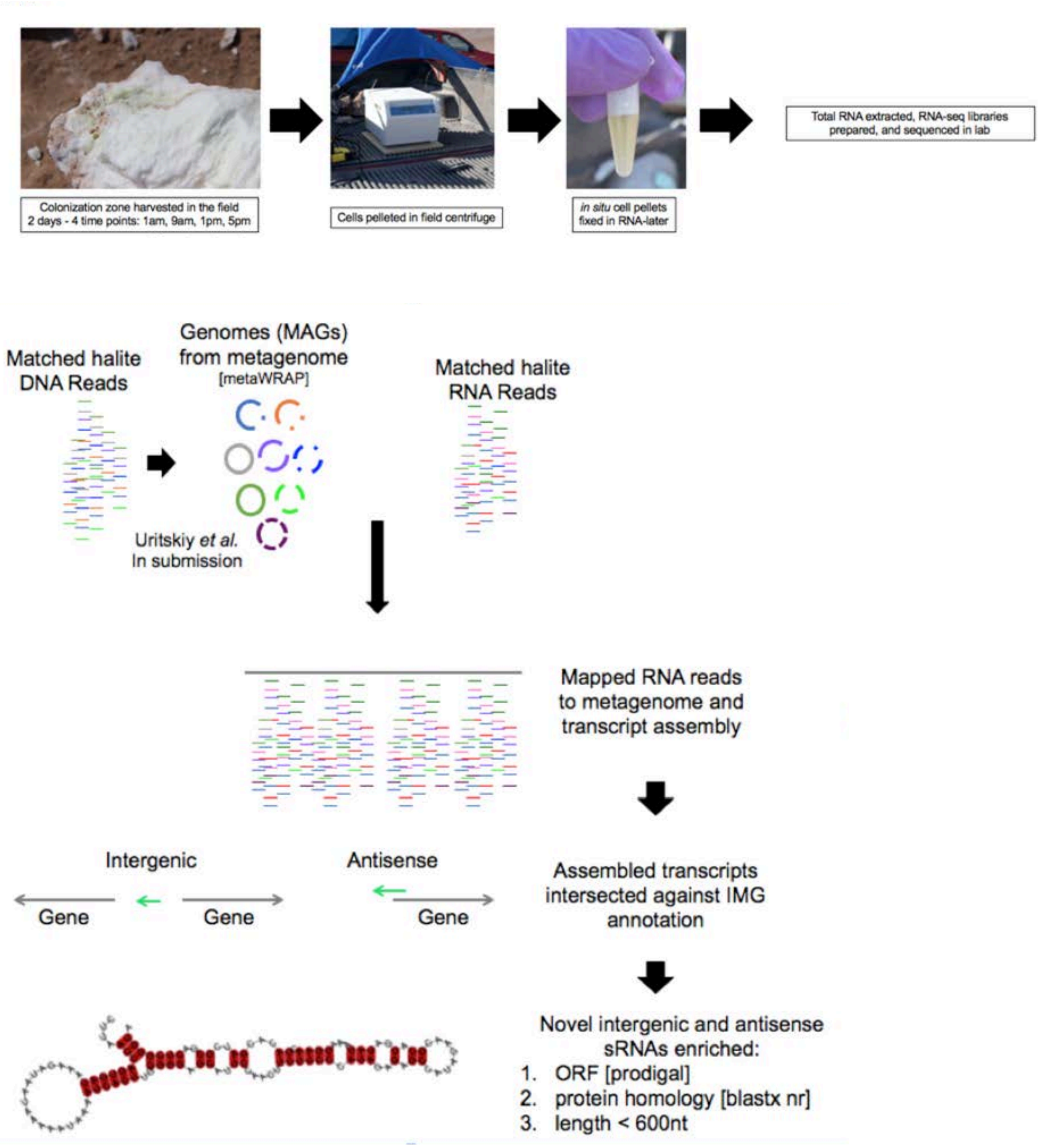


**Fig. S1** Experimental design

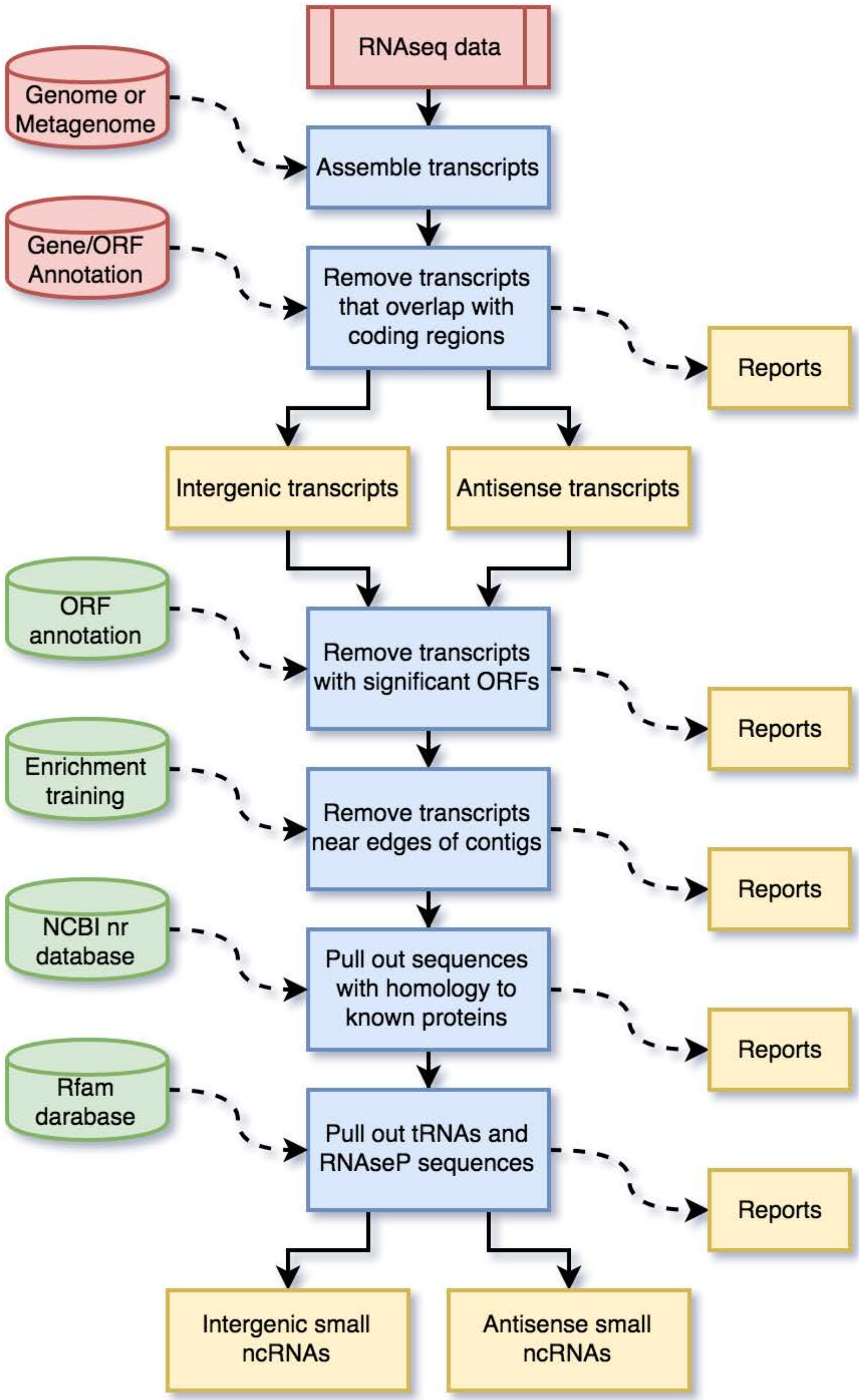


**Fig. S2** Flow chart for SnapT methodology

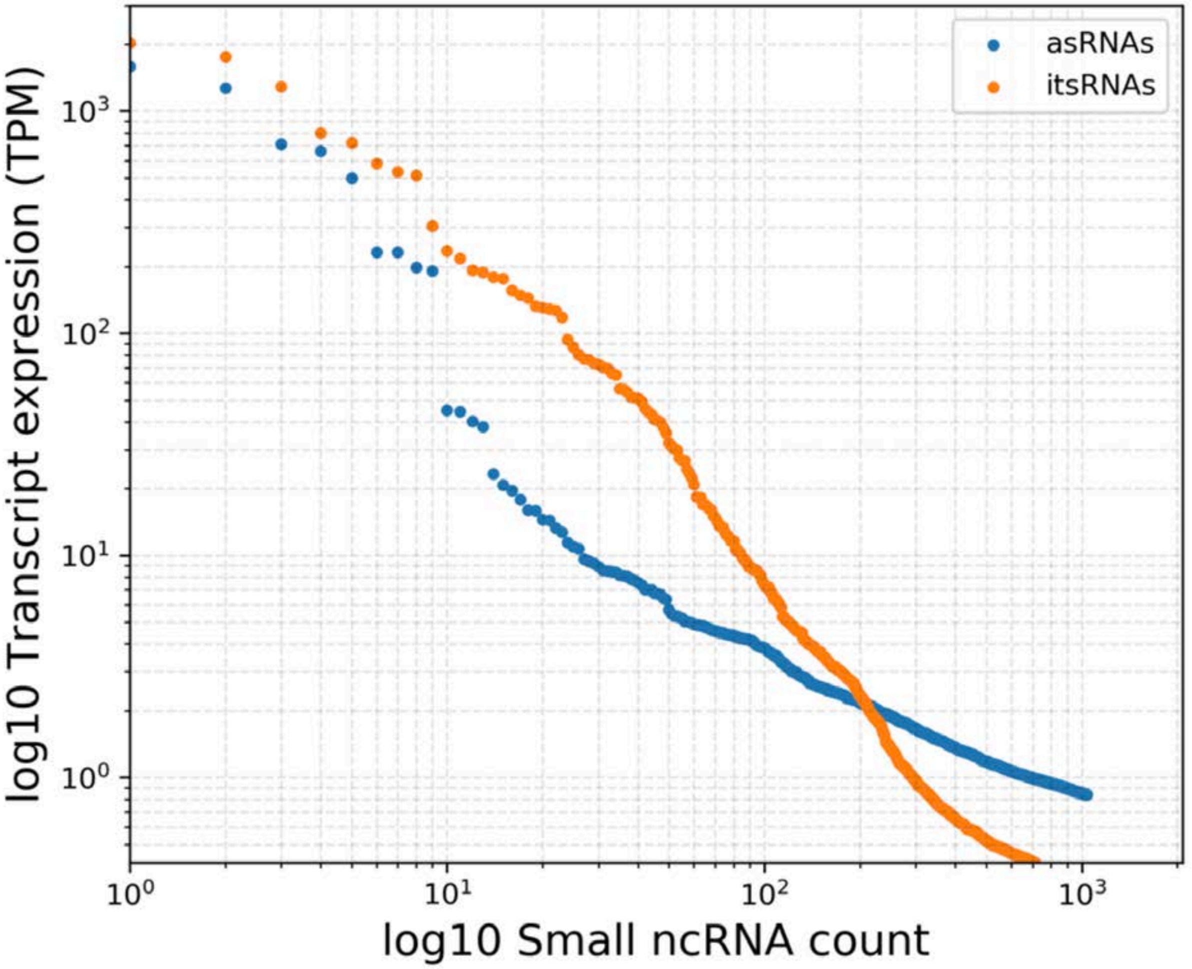


**Fig. S3** Thresholding for its and asRNAs. Ranked total expression (transcripts per million) of annotated antisense and intergenic small ncRNAs. The figure shows the linear relationship between the sRNAs expression in TPM and the number of sRNAs and how it decays. A threshold at 5x and 10x coverage was applied to its and asRNAs, respectively.

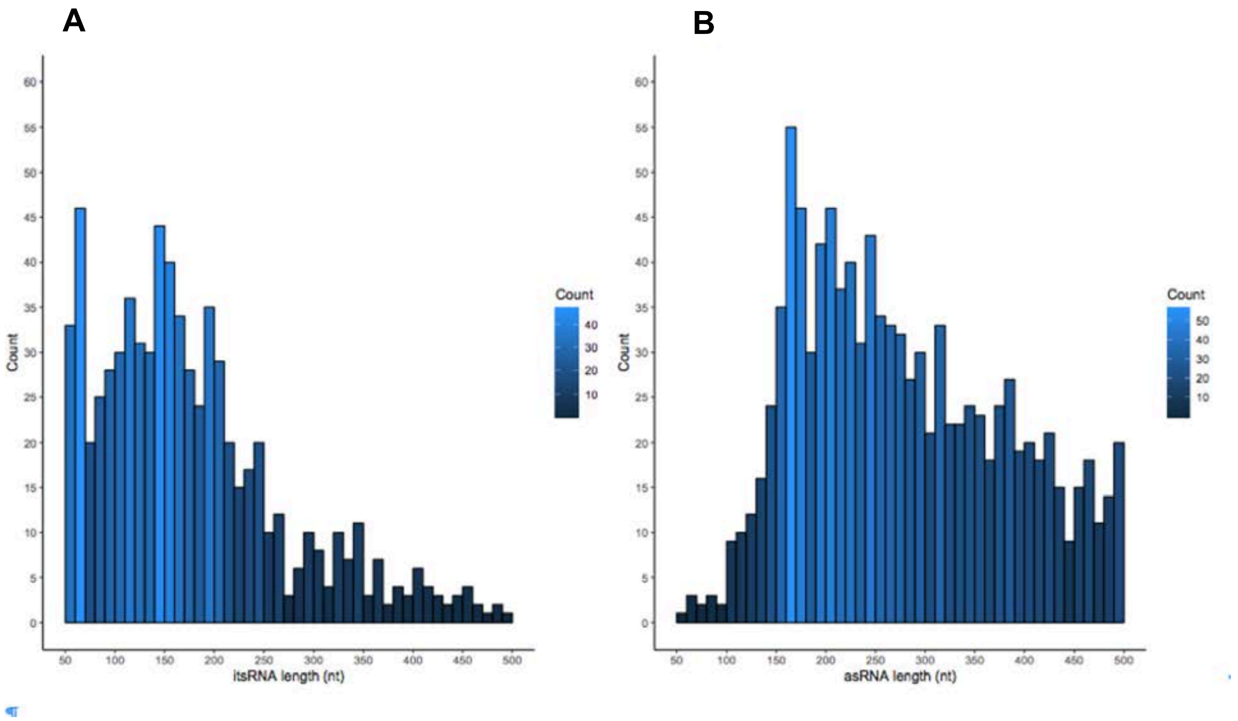


**Fig. S4** Length distribution of (a) itsRNAs and (B) asRNAs

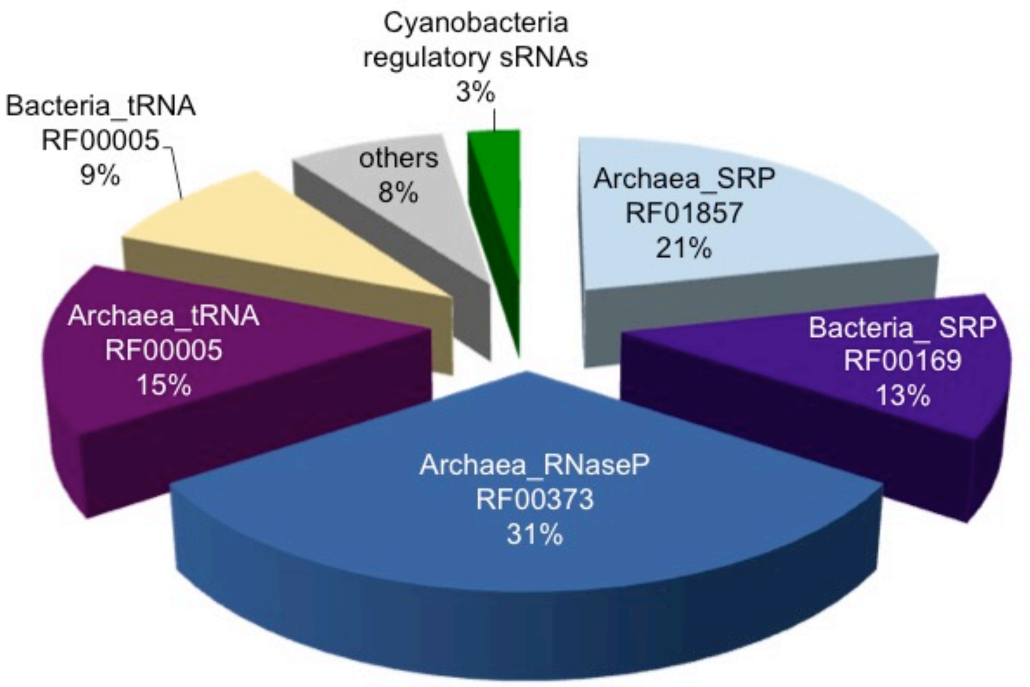


**Fig. S5** Rfam-conserved sRNAs indentified the halite community

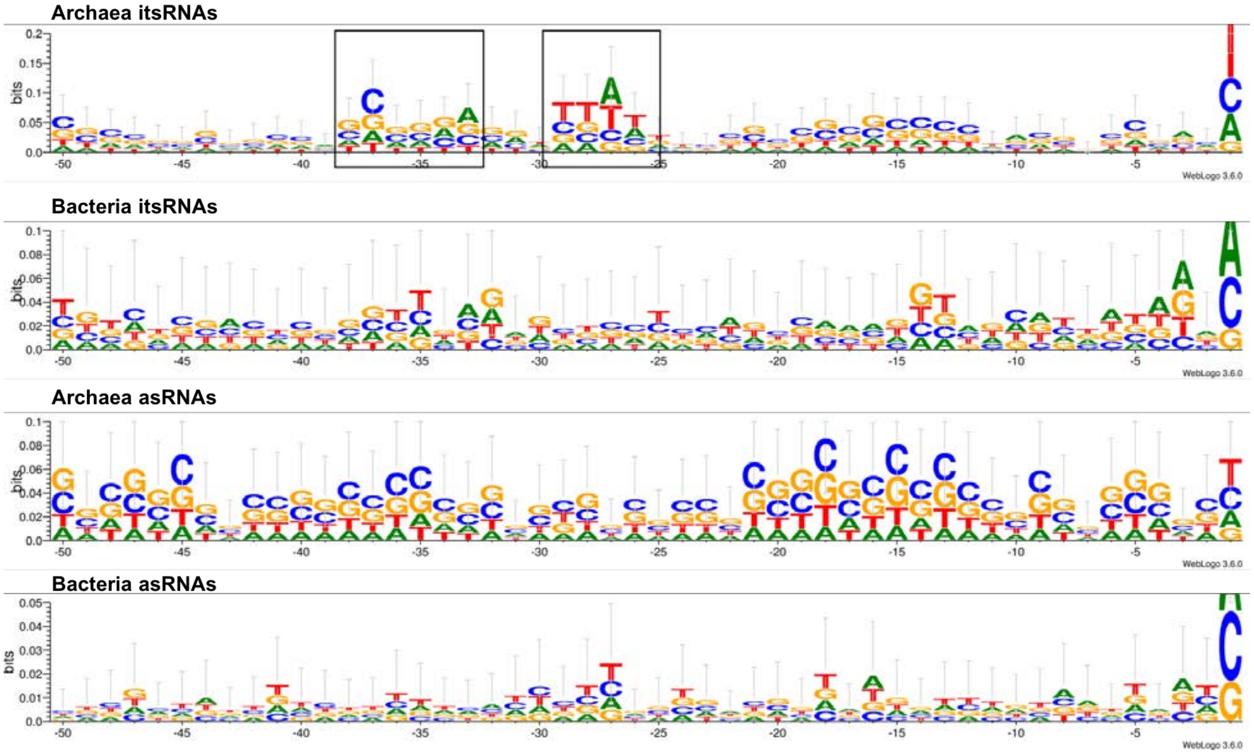


**Fig. S6** Regulatory regions for asRNAs and itsRNAs and for archaea and bacteria

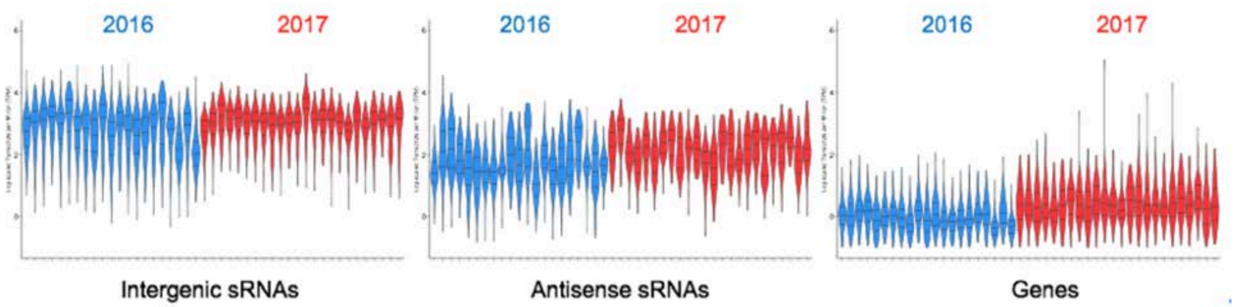


**Fig. S7** Expression level of itsRNAs, asRNAs, and protein encoding genes normalized by contig abundance

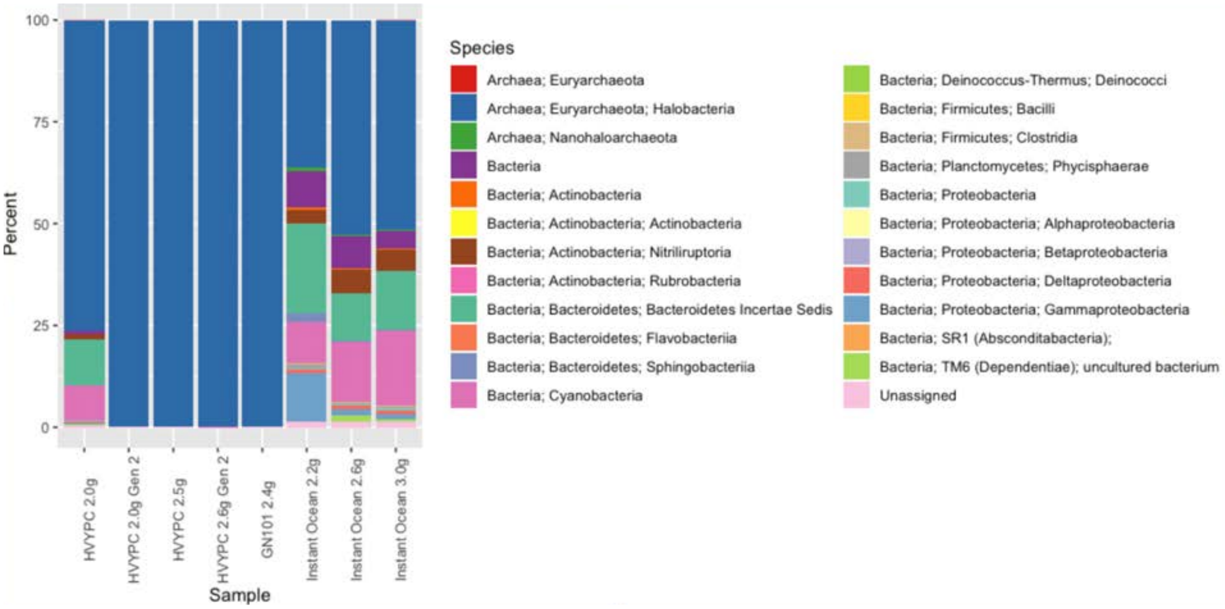


**Fig. S8** Taxonomy distribution of halite enrichment cultures showing the relative abundance (%) of bacterial and archaeal taxa in function of the culture media.

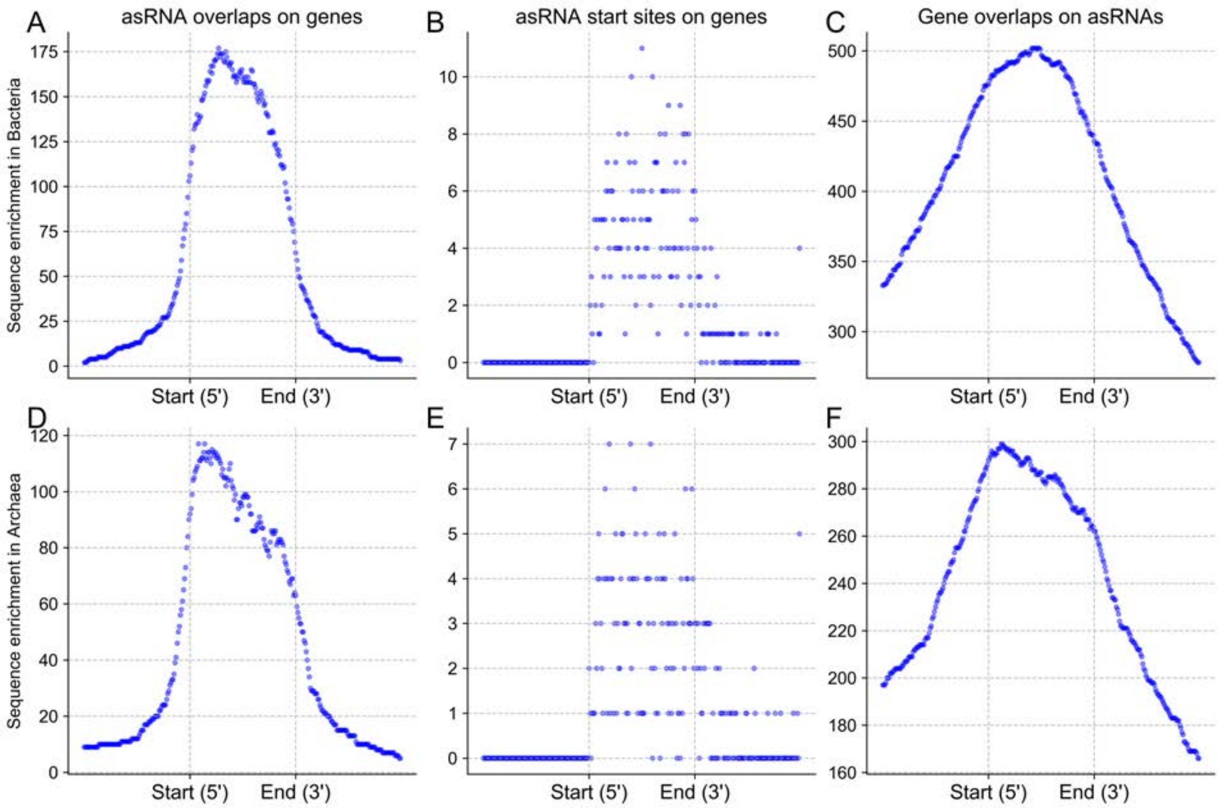


**Fig. S9** asRNA overlap distribution with their putative mRNAs

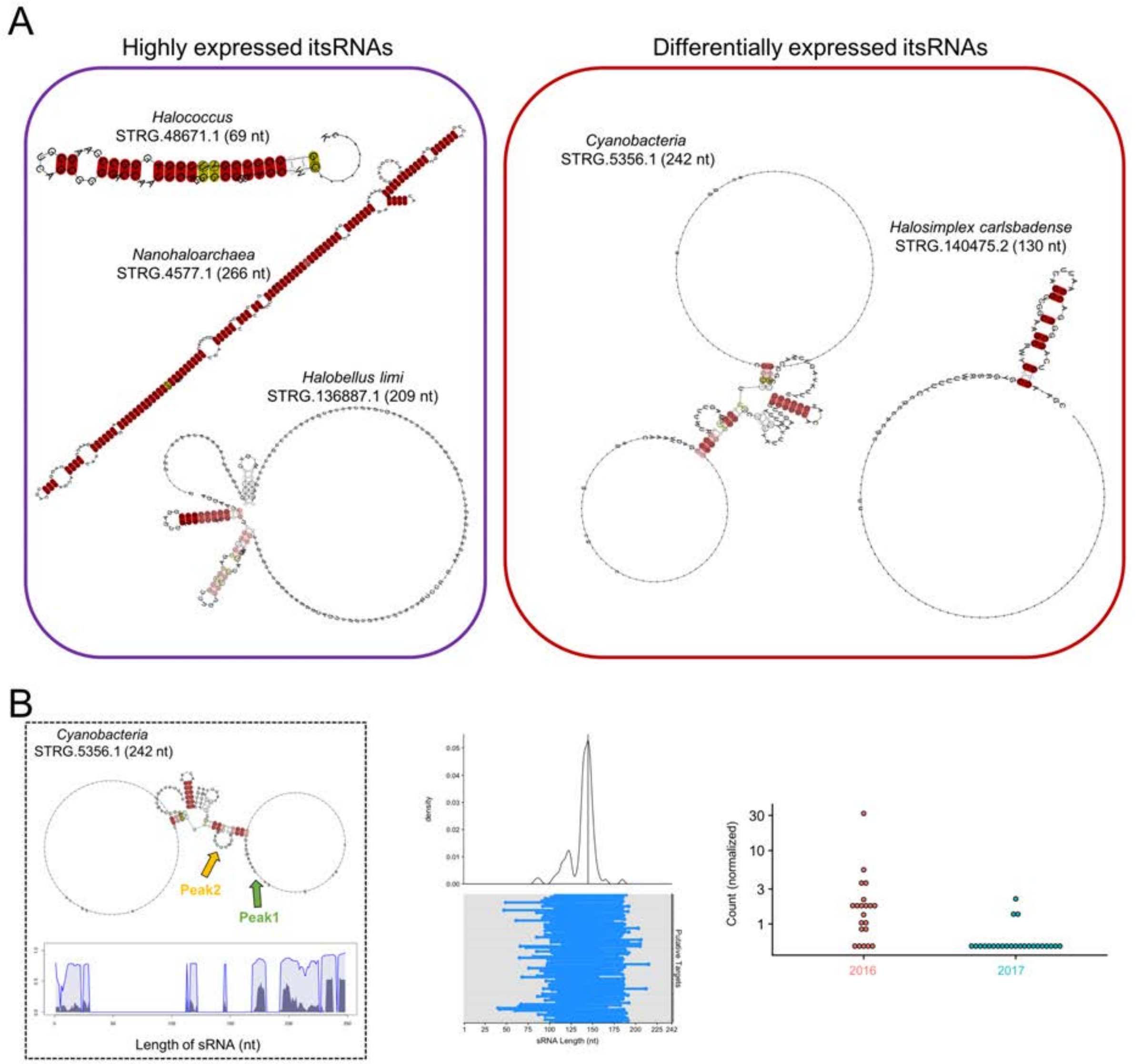


**Fig. S10** (A) Additional structure prediction for highly expressed and differential expressed itsRNAs; (B) Predicted structure, target identification, and expression levels for differentially expressed itsRNA STRG.5356.1 from a cyanobacterium.

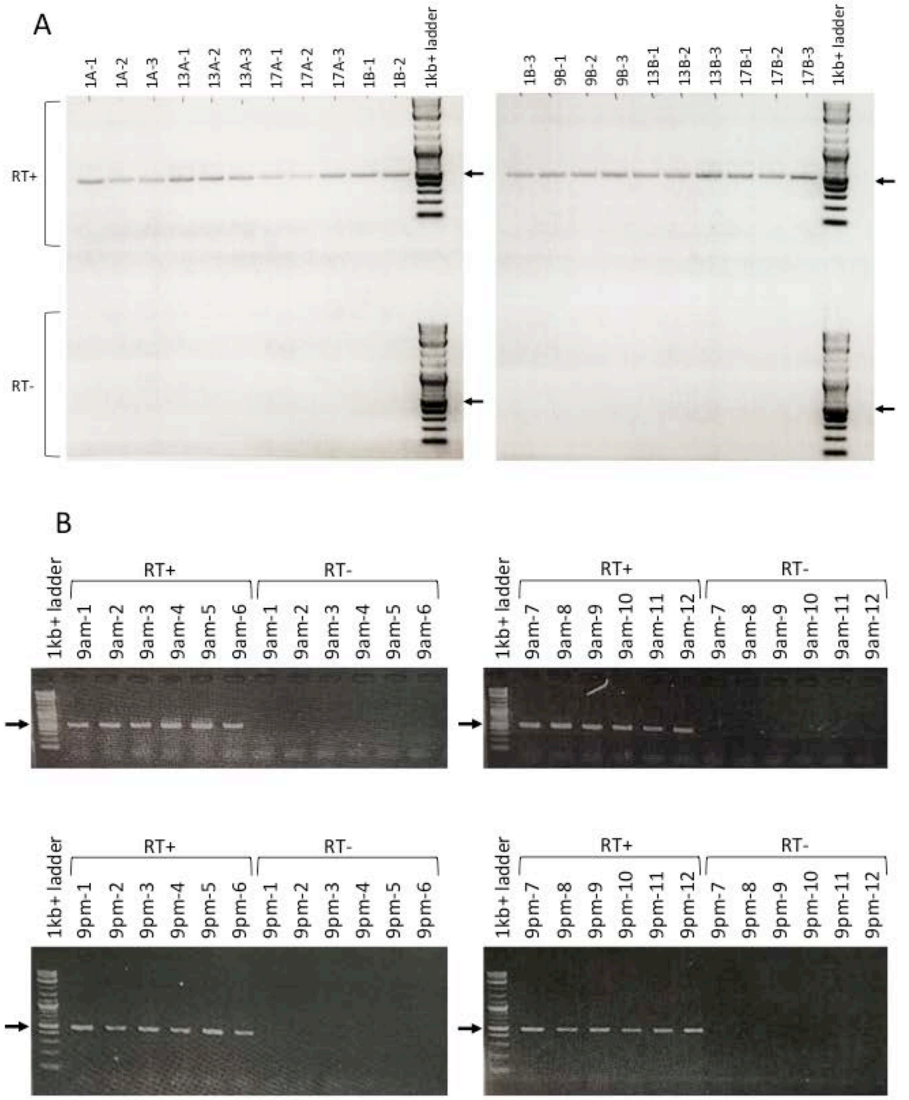


**Fig. S11** RT-PCR validation showing the absence of DNA contamination in the total RNA used for RNA-seq library preparation. (A) Samples collected in 2017 and (B) samples collected in 2016. Each RNA sample was processed using the SuperScript III cDNA synthesis kit with (RT+, positive control) or without (RT-, negative control) the reverse transcriptase enzyme. The 16S rRNA gene was amplified from the resulting cDNAs. Arrows indicate the expected locations of amplicon bands (512bp).
